# Molecular basis for SPIN·DOC-Spindlin1 engagement and its role in transcriptional inhibition

**DOI:** 10.1101/2021.03.07.432812

**Authors:** Fan Zhao, Fen Yang, Fan Feng, Bo Peng, Mark T. Bedford, Haitao Li

## Abstract

Spindlin1 is a transcriptional coactivator with three Tudor-like domains, of which the first and second Tudors are engaged in histone methylation readout, while the function of the third Tudor is largely unknown. Recent studies revealed that the transcriptional co-activator activity of Spindlin1 could be attenuated by SPIN•DOC. Here we solved the crystal structure of SPIN•DOC-Spindlin1 complex, revealing that a hydrophobic motif, DOCpep3 (256-281), of SPIN•DOC interacts with Tudor 3 of Spindlin1 and completes its β-barrel fold. Massive hydrophobic contacts and hydrogen bonding interactions ensure a high affinity DOCpep3-Spindlin1 engagement with a binding *K*_d_ of 30 nM. Interestingly, we characterized two more K/R-rich motifs of SPIN•DOC, DOCpep1 (187-195) and DOCpep2 (228-239), which bind to Spindlin1 at lower affinities with *K*_d_ values of 78 μM and 31 μM, respectively. Structural and binding studies revealed that DOCpep1 and DOCpep2 competitively bind to the aromatic cage of Spindlin1 Tudor 2 that is responsible for H3K4me3 readout. Although DOCpep3-Spindlin1 engagement is compatible with histone readout, an extended SPIN•DOC fragment containing DOCpep1 and DOCpep2 inhibits histone or TCF4 binding by Spindin1 due to introduced competition. This inhibitory effect is more pronounced for weaker binding targets but not for strong ones such as H3 “K4me3-K9me3” bivalent mark. Our RT-qPCR experiment showed that the removal of the hydrophobic motif or the K/R-rich region compromised the inhibitory effects of SPIN•DOC on Spindlin1-mediated transcriptional activation. In sum, here we revealed multivalent engagement between SPIN•DOC and Spindlin1, in which a hydrophobic motif acts as the primary binding site for stable SPIN•DOC-Spindlin1 association, while two more neighboring K/R-rich motifs further modulate the target selectivity of Spindlin1 via competitive inhibition, therefore attenuating the transcriptional co-activator activities of Spindlin1 through affecting its chromatin association.

## INTRODUCTION

As the physiological carrier of genetic information in eukaryotic cells, chromatin provides an epigenetic layer of regulation for gene expression. In particular, the basic and flexible histone tails could undergo diverse post-translational modifications, among which include methylation, acetylation, phosphorylation as well as lactylation, benzoylation and serotonylation that were discovered recently (1–4). These modifications often serve as a docking mark for downstream “reader” modules to facilitate gene regulation and other chromatin-templated processes (2,5). Histone methylation mainly occurs on arginine or lysine residues scattered at different sites of histones. These methylation marks may act alone or in combination to demarcate distinct gene expression states (e.g. “active”, “repressive” or “poised”) (6,7).

Spindlin1 contains three tandem Tudor-like domains and has been characterized as a reader for multiple histone methylation patterns (8). For example, Spindlin1 recognizes H3K4me3 or H3 “K4me3-R8me2a” methylation pattern to activate expression of rDNA genes, Wnt/TCF4 or MAZ (Myc-associated zinc finger protein) target genes (9–14). Spindlin1 also recognizes H4K20me3, a marker in silenced heterochromatin regions, which might regulate DNA replication and cell cycle progression (15). We recently revealed that Spindlin1 can serve as a potent H3 “K4me3-K9me3/2” bivalent histone methylation reader to balance gene expression and silencing in H3K9me3/2-enriched regions (7). Intriguingly, Spindlin1 can also associate with spindle and play a role in spindle stability maintenance and chromosome segregation during cell division (16–21). Spindlin1 is highly expressed in multiple cancers including colorectal cancer, cervical cancer and liposarcoma, suggesting its tumor promoter role (12, 14, 22–27). Among the three Tudor-like domains, it has been well established that the first and second Tudors participate in histone methylation readout, while the function of the third Tudor is largely unknown (7,9,10,13,15).

SPIN•DOC (SPIN1 docking protein) has been identified as a Spindlin1-interacting protein and negatively regulates Spindlin1’s transcriptional co-activator activity (28,29). However, the molecular basis for SPIN•DOC-Spindlin1 engagement and the resultant transcriptional inhibition remain to be explored. Here we performed domain mapping, crystal structural determination and cell-based functional validation studies. Our structural and binding studies revealed two low affinity K/R-rich motifs (DOCpep1 and DOCpep2) and one high affinity hydrophobic motif (DOCpep3) that are responsible for Spindlin1 interaction. In details, DOCpep3 binds to Tudor 3 of Spindlin1 at nanomolar level affinity and completes its β-barrel fold. Meanwhile, DOCpep1 and DOCpep2 binds to the histone binding Tudor 2 surface of Spindlin1 at micromolar level affinities. We further showed that although DOCpep3-Spindlin1 engagement is compatible with histone readout, the existence of neighboring DOCpep1 and DOCpep2 motifs inhibits Spindlin1’s histone or other ligand binding that requires an available Tudor 2 surface. Our RT-PCR experiments validated a role of both DOCpep1/2 and DOCpep3 motifs of SPIN•DOC in attenuating the co-activator activity of Spindlin1. Collectively, our study provides molecular insights into SPIN•DOC inhibitory function that involves in mechanisms of both multivalent engagement and competitive inhibition.

## MATERIALS AND METHODS

### Protein and peptides preparation

To obtain SPIN•DOC-Spindlin1 complex, the full-length genes of SPIN•DOC and Spindlin1 were cloned into the first and second multiple cloning sites of pRSFDuet vector, respectively. The N terminal of SPIN•DOC has a 6xHis tag, while Spindlin1 has no tag. Spindlin1 and SPIN•DOC genes were transformed to Escherichia coli BL21 (DE3) strain for overexpression. After induced by 0.2 mM isopropyl β-D-thiogalactoside (IPTG) for 12-18 hours at 16°C in LB medium, the cells were collected and suspended in buffer containing 200 mM NaCl, 20 mM Tris, pH 8.0. After cell lysis and centrifugation, the supernatant was applied to a HisTrap column (GE Healthcare) for affinity purification followed by anion-exchange chromatography over a HiTrap Q HP column. Then, the resultant protein was further polished by a Superdex 200 gel filtration column (GE Healthcare). The purified protein was concentrated up to 60 mg/ml in buffer containing 150 mM NaCl, 20 mM Tris, pH 8.0, and aliquoted, stored at −80°C for future use.

Human Spindlin1_50-262_ was cloned into pGEX-6p vector containing an N-terminal GST tag. Recombinant protein was overexpressed and purified using essentially the same procedures as described above. Briefly, after cell lysis and centrifugation, the supernatant was purified by a GST affinity column. The GST tag was cleaved on-column by a home-made P3C protease overnight. The tag-free protein was collected for an anion-exchange chromatography over a HiTrap Q HP column and then polished over a Superdex 75 gel filtration column (GE Healthcare). Peak fractions were pooled and concentrated to 25 mg/ml in buffer containing 150 mM NaCl, 20 mM Tris, pH 8.0, and aliquoted, stored at −80°C for future use.

All Spindlin1 mutants were purified using essentially the same procedures as described above. All synthetic histone and SPIN•DOC peptides (>95% purity) were synthesized by and purchased from Scilight Biotechnology LLC.

### Isothermal titration calorimetry

Isothermal titration calorimetric experiments were conducted with a MicroCal PEAQ-ITC instrument (Malvern) at 25°C. The Spindlin1_50-262_ sample and peptides were prepared in ITC buffer containing 150 mM NaCl, 20 mM HepesNa, pH 7.5. Protein concentration was measured by UV absorption at 280 nm, while peptides were quantified by weighing in a large scale. About 200 μl protein sample in cell was titrated with the corresponding peptide in syringe. Titration curves were fitted using the “One-Set-of-Binding-Sites” model with the Origin 7.0 program.

### Surface plasmon resonance

Peptide of SPIN•DOC 256-281 was immobilized onto the SPRi (SPR imaging) chip surface using an EDC/NHS coupling strategy. Wild type or mutant Spindlin1 samples in different concentrations were applied to the chip surface. A Kx5 instrument (Plexera, USA) was used for signal monitor. The binding signals were recorded for data analyses using the commercial software (Plexera SPR Data Analysis Module, Plexera, USA).

### X-ray crystallographic studies

The full length SPIN•DOC-Spindlin1 complex was mixed with trace amount of trypsin protease at a ratio of 1:5000 (w/w) for in situ proteolysis crystallization. The mixture sample was applied for crystallization screen by mixing 200 nl SPIN•DOC-Spindlin1 complex with 200 nl reservoir solution from the commercial crystallization kits. Crystals were obtained under the buffer condition: 20% PEG8000, 0.2 M magnesium acetate and 0.1 M Sodium cacodylate, pH 6.5.

Spindlin1_50-262_ and DOCpep2 were mixed at 1:5 ratio and incubated at 4°C for 1 hour. The sample was subjected to crystallization screen by mixing 200 nl of protein sample with 200 nl of reservoir solution. The crystal was obtained in a buffer condition: 1.5 M Ammonium Sulfate, 0.1 M BIS-TRIS, pH 6.5, and 0.1 M Sodium chloride.

Crystals were flash-frozen with supplemented 30% (v/v) glycerol cryo-protectant in liquid nitrogen for further data collection at beamline BL17U of Shanghai Synchrotron Radiation Facility. Diffraction data sets were indexed, integrated and merged using the HKL2000 suite (http://www.hkl-xray.com). Crystal structures were determined by molecular replacement using Spindlin1 structure (PDB code 2NS2) as a search model. Model building and refinement were performed using software COOT (30) and PHENIX (31), respectively. Data processing and structural refinement statistics are summarized in Table 1.

**Table 1.**
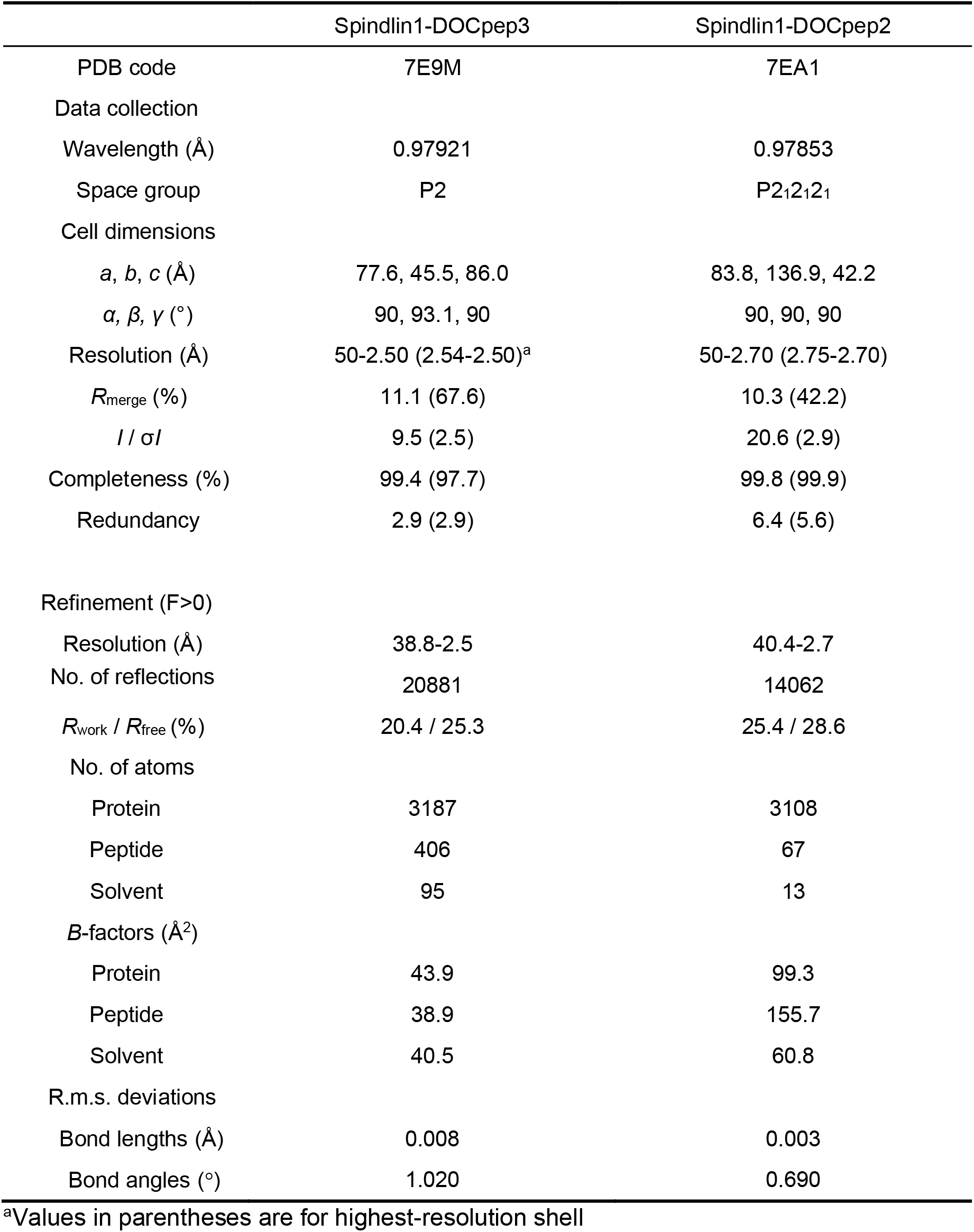
Data collection and refinement statistics.

|  | Spindlin1-DOCpep3 | Spindlin1-DOCpep2 |
| --- | --- | --- |
| PDB code | 7E9M | 7EA1 |
| Data collection |  |  |
| Wavelength (Å) | 0.97921 | 0.97853 |
| Space group | P2 | P2<sub>1</sub>2<sub>1</sub>2<sub>1</sub> |
| Cell dimensions |  |  |
| *a*, *b*, *c* (Å) | 77.6, 45.5, 86.0 | 83.8, 136.9, 42.2 |
| $\alpha$, $\beta$, $\gamma$ (°) | 90, 93.1, 90 | 90, 90, 90 |
| Resolution (Å) | 50-2.50 (2.54-2.50)<sup>a</sup> | 50-2.70 (2.75-2.70) |
| *R*<sub>merge</sub> (%) | 11.1 (67.6) | 10.3 (42.2) |
| *I* / σ*I* | 9.5 (2.5) | 20.6 (2.9) |
| Completeness (%) | 99.4 (97.7) | 99.8 (99.9) |
| Redundancy | 2.9 (2.9) | 6.4 (5.6) |
| Refinement (*F*>0) |  |  |
| Resolution (Å) | 38.8-2.5 | 40.4-2.7 |
| No. of reflections | 20881 | 14062 |
| *R*<sub>work</sub> / *R*<sub>free</sub> (%) | 20.4 / 25.3 | 25.4 / 28.6 |
| No. of atoms |  |  |
| Protein | 3187 | 3108 |
| Peptide | 406 | 67 |
| Solvent | 95 | 13 |
| *B*-factors (Å<sup>2</sup>) |  |  |
| Protein | 43.9 | 99.3 |
| Peptide | 38.9 | 155.7 |
| Solvent | 40.5 | 60.8 |
| R.m.s. deviations |  |  |
| Bond lengths (Å) | 0.008 | 0.003 |
| Bond angles (°) | 1.020 | 0.690 |

<sup>a</sup>Values in parentheses are for highest-resolution shell

### Cell culture and recombinant plasmids transfection

HEK293T and T778 cell lines were purchased from the ATCC. All cell lines used in this study were routinely tested for mycoplasma by using MycoAlert™ (Lonza). T778 cell line was cultured in RPMI 1640 medium supplemented with 10% FBS, 1% penicillin/streptomycin and L-glutamine. HEK293T cell line was cultured in DMEM supplemented with 10% FBS, 1% penicillin/streptomycin, nonessential amino acids and L-glutamine. All cells were maintained in a humidified 37 °C incubator with 5% CO2. HEK293 cells were transfected or co-transfected with vector controls, Flag-SPIN1, GFP-SPIN•DOC and corresponding mutants using polyethyleneimine (PEI) according to the instructions of the manufacturer for 48 hours. Transfected whole cell lysates were immunoprecipitated and subjected to Western blot analysis. T778 cells were transfected or co-transfected with vector controls, GFP-SPIN1, Flag-SPIN•DOC and Flag-SPIN•DOC mutants using PEI for 48 hours.

### Co-Immunoprecipitation (Co-IP) and Western blot analysis

HEK293 cells were lysed with RIPA lysis buffer containing protease inhibitor cocktail and DNase I on ice for 30 mins and centrifuged with 15000 rpm for 10 mins. The supernatant was incubated with 2 μg primary antibody overnight rotation at 4°C and added with 25 μl protein A/G agarose beads for an hour rotation at 4°C. Beads were washed and boiled in 2xSDS buffer and the protein separated by SDS-PAGE, and then transferred onto a PVDF membrane and then blocked in 5% non-fat milk in PBST (0.05% Tween 20 with 1xPBS) buffer. Subsequently, the PVDF membranes were probed with the indicated primary and secondary antibodies, and signal was detected using AmershamTM ECL detection reagents (GE Healthcare).

### GST fusion protein purification and GST pulldowns

For the GST pulldown assay, we inoculated a single plasmid-containing bacterial colony into 100 ml of fresh LB Broth with antibiotic and incubate for an hour, shaking at 37°C. IPTG (Sigma) was then added to final concentration of 0.1 mM and incubated for 4 hours, shaking at 30°C. Cells were then centrifuged and the pellets resuspended in 500 μl of ice-cold PBS. Samples were then sonicated at 30% amplitude using pulsations (0.5s on, 0.5s off) for 20s. After centrifugation, 100 μl Glutathione sepharose beads (GE LifeScience) were added to the supernatant, rotated overnight at 4°C, and washed beads 3 times with ice-cold 1x PBS. We then added purified His-SPIN•DOC_251-293_ protein or cell lysates from 293T cells transfected with GFP-SPIN•DOC_251-293_, and rotated overnight at 4°C. Beads were then washed and boiled in 2xSDS buffer. Finally, Western blot was performed with anti-GST, -His and -GFP antibodies.

For His-fusion protein purification, the initial steps were the same as above, but samples were lysed in the special lysis buffer (50 mM NaH_2_PO_4_, 300 mM NaCl, 10 mM imidazole, pH 8.0) and nickel beads were used. After incubation overnight at 4°C, the beads were washed 3 times with wash buffer (50 mM NaH_2_PO_4_, 300 mM NaCl, 20 mM imidazole, pH 8.0), and then eluted with elution buffer (50 mM NaH_2_PO_4_, 300 mM NaCl, 200 mM imidazole, pH 8.0) 200 μl with rotation overnight at 4°C. After centrifugation the supernatant harbored the purified fusion protein.

### Quantitative real-time PCR (RT-qPCR)

Total RNA was extracted from T778 cells using the RNeasy Mini Kit (Qiagen) and reverse-transcribed using a Superscript III First Strand Synthesis Kit (Invitrogen). QPCR was performed with the Applied Biosystems 7900HT RT-PCR instrument using SYBR Green Supermix (Bio-Rad) with indicated primers. All primers used in this study were synthesized by Integrated DNA Technologies (IDT) and gene expression was calculated following normalization to GAPDH levels using the comparative cycle threshold method and was shown as folds relative to the expression of each gene in the control cells. The primer sequences were as follows. SPIN1: Forward, ATGAAGACCCCATTCGGAAA and Reverse, TGTGGGATGTCCTCTTCTTC. SPIN•DOC: Forward, CAGGGAAGTGGCAGAAGGAG and Reverse, CACGCGGATAACCTGGAGAT. IFI44L: Forward, CCGTCAGTATTTGGAATGTGAAG and Reverse, TGAAACC AAGTCTGCATAGGG. BST2: Forward, TGATGGAGTGTCGCAATGTC and Reverse, GTCCTTGGGCCTTCTCTG. IL1B: Forward, GAGGGAGAAACTGGCAGATACC and Reverse, TCTGTTTAGGGCCATCAGCTT. CyclinD1: Forward, CCCTCGGTGTCCTACTT CAAA and Reverse, GAAGACCTCCTCCTCGCACT. β-actin: Forward, CTTCGCGGGCGACGAT and Reverse, CCACATAGGAATCCTTCTGACC. GAPDH: Forward, AGCCACATCGCTCAGACAC and Reverse, GCCCAATACGACCAAATCC.

## RESULTS

### Spindlin1 stably interacts with SPIN•DOC via its Tudor 3 domain

Spindlin1 is 262 amino acids in length and is composed of an N-terminal flexible tail and three Tudor-like repeats (Figure 1A). The first and second Tudor domains are responsible for histone methylation readout, while the function of the third Tudor domain remains largely unknown (7,9,10,13,15). Full-length SPIN•DOC has 381 residues, most of which are flexible regions, and its middle fragment (184-240) is characteristic of K/R-rich motifs followed by a hydrophobic region (Figure 1A, left). To elucidate the molecular basis for SPIN•DOC-Spindlin1 interaction, we co-expressed full-length SPIN•DOC and Spindlin1 and successfully reconstituted the binary complex (Figure 1A, right). Then we performed in situ proteolysis crystallization screening by mixing the full length binary complex with trace amount (1/5000 w/w) of trypsin protease. We obtained diffractable crystals through this strategy and finally solved a complex structure at 2.5 Å resolution (Table 1). In the crystal structure, a “β1-loop-β2” motif of SPIN•DOC binds to Tudor 3 of Spindlin1 and completed its β-barrel fold (Figure 1B). According to the electron density map, we were able to assign a hydrophobic fragment 256-281 of SPIN•DOC (DOCpep3) to the “β1-loop-β2” motif (Figure 1C). Electrostatic surface analysis revealed that DOCpep3 binds to a hydrophobic surface of Spindlin1 with sound shape complementarity (Figure 1D). Using a surface plasmon resonance (SPR) instrument, we could measure a dissociation constant (*K*_d_) of 30 nM between DOCpep3 and Spindlin1_50-262_ (Figure 1E), underlining their strong association.

**Figure1.**
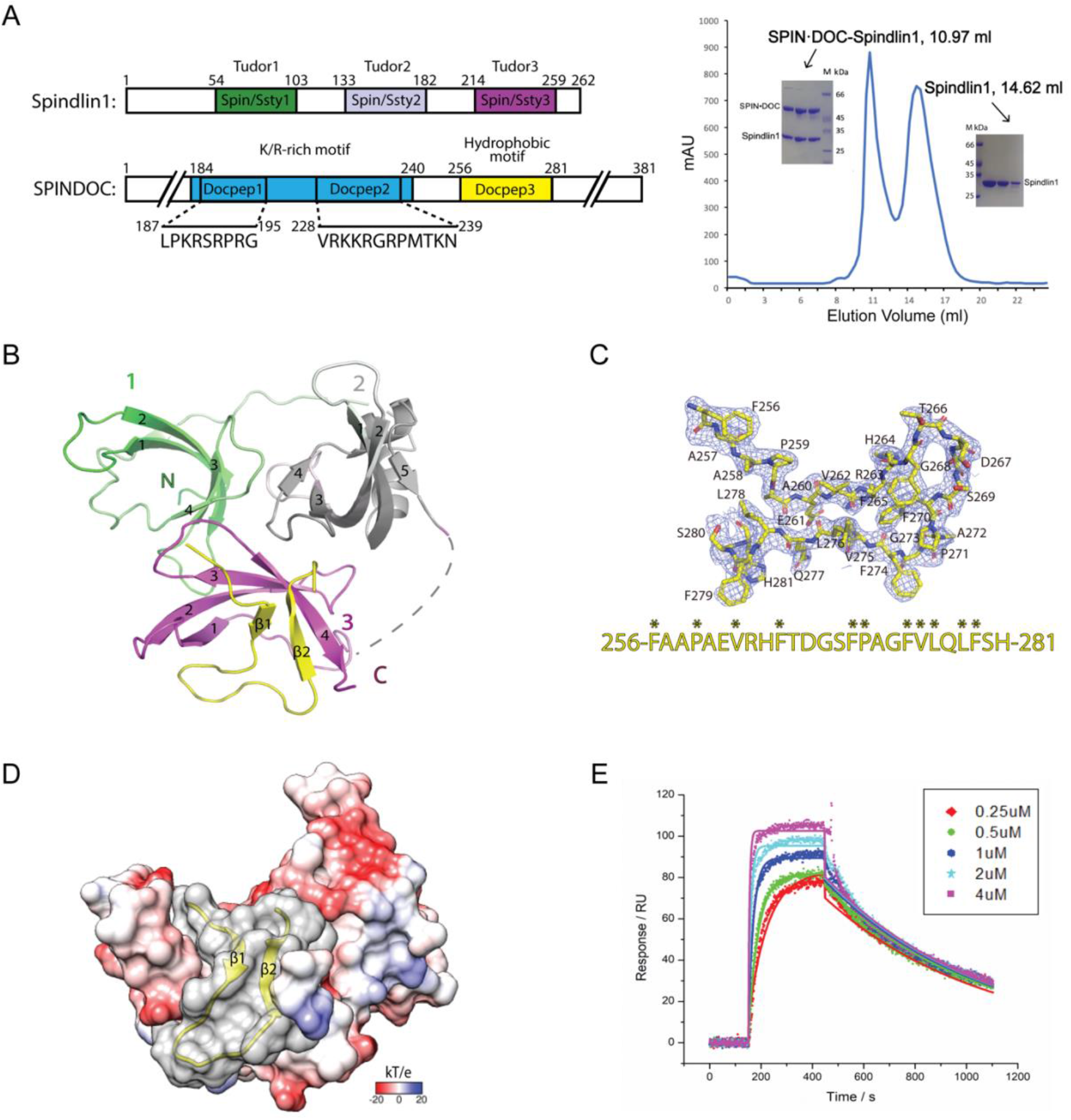
Crystallographic characterization of SPIN•DOC as a Spindlin1 binding partner. **(A)** (left) Domain architecture of Spindlin1 and SPIN•DOC. The core β-barrel regions of each Spin/Ssty repeat in Spindlin1 are colored green, white, and magenta, respectively. (Black line) Construct Spindlin1_50–262_ used for ITC and structural studies. In SPIN•DOC, K/R-rich region (DOCpep1-2) and DOCpep3 are indicated as blue and yellow box. (right) Size exclusion chromatography of full length SPIN•DOC-Spindlin1 complex and SDS-PAGE gel of the complex and Spindlin1 alone. **(B)** Overall structure of Spindlin1 bound to DOCpep3 in ribbon view. DOCpep3 is depicted as yellow ribbon. **(C)** 2Fo-Fc omit map of DOCpep3 contoured at the 1.5 σ level and its amino acid sequence (hydrophobic residue labeled by asterisk) are shown here. **(D)** Electrostatic potential surface view of Spindlin1-DOCpep3 complex. Electrostatic potential is expressed as a spectrum ranging from −20 kT/e (red) to +20 kT/e (blue). **(E)** SPR fitting curves of five gradients of Spindlin1_50–262_ concentration flowing immobilized DOCpep3.

### Details of Spindlin1-DOCpep3 interaction and mutagenesis studies

In the complex structure, DOCpep3 interacts with Spindlin1 via extensive hydrophobic interactions involving residues V262, L276, F265 and F270 that are clustered in the center core and residues F256, F274, F279 that are anchored at three corners (Figure 2A). The stable engagement between DOCpep3 and Spindlin1 is also warranted by backbone-mediated hydrogen bonding interactions, including an anti-parallel “β1-β2-β4_(Tudor3)_” sheet and a pseudo-β-sheet between the “β1-loop” and “β1_(Tudor 3)_” (Figure 2B). Additionally, the binding is further contributed by a couple of side-chain mediated hydrogen bonds involving the “K216-Q217” step of Tudor 3 and the “T266-D267-S69” segment of DOCpep3 (Figure 2B). Three hydrophobic residues, V218, V232, and I245 of Tudor 3 constitute the hydrophobic core with V262, F265, F270 and L276 of DOCpep3 (Figure 2C). Next, we generated V218R, V232R and I245R single point mutants and performed SPR binding assays. All three mutants were well expressed and highly purified, yet they displayed nearly no binding towards immobilized DOCpep3 peptide as compared to the wild type Spindlin1, stressing the importance of these hydrophobic residues (Figure 2D). Structural alignment revealed conformational adjustments between the free-state and DOCpep3-bound Tudor 3 of Splindlin1, in which the “β1-loop-β2” hairpin of Tudor 3 was straighten up upon DOCpep3 binding, notably trigged by the insertion of F256 at the very N-terminus of DOCpep3 and the resultant displacement of R228 of Tudor 3 (Figure 2E).

**Figure 2.**
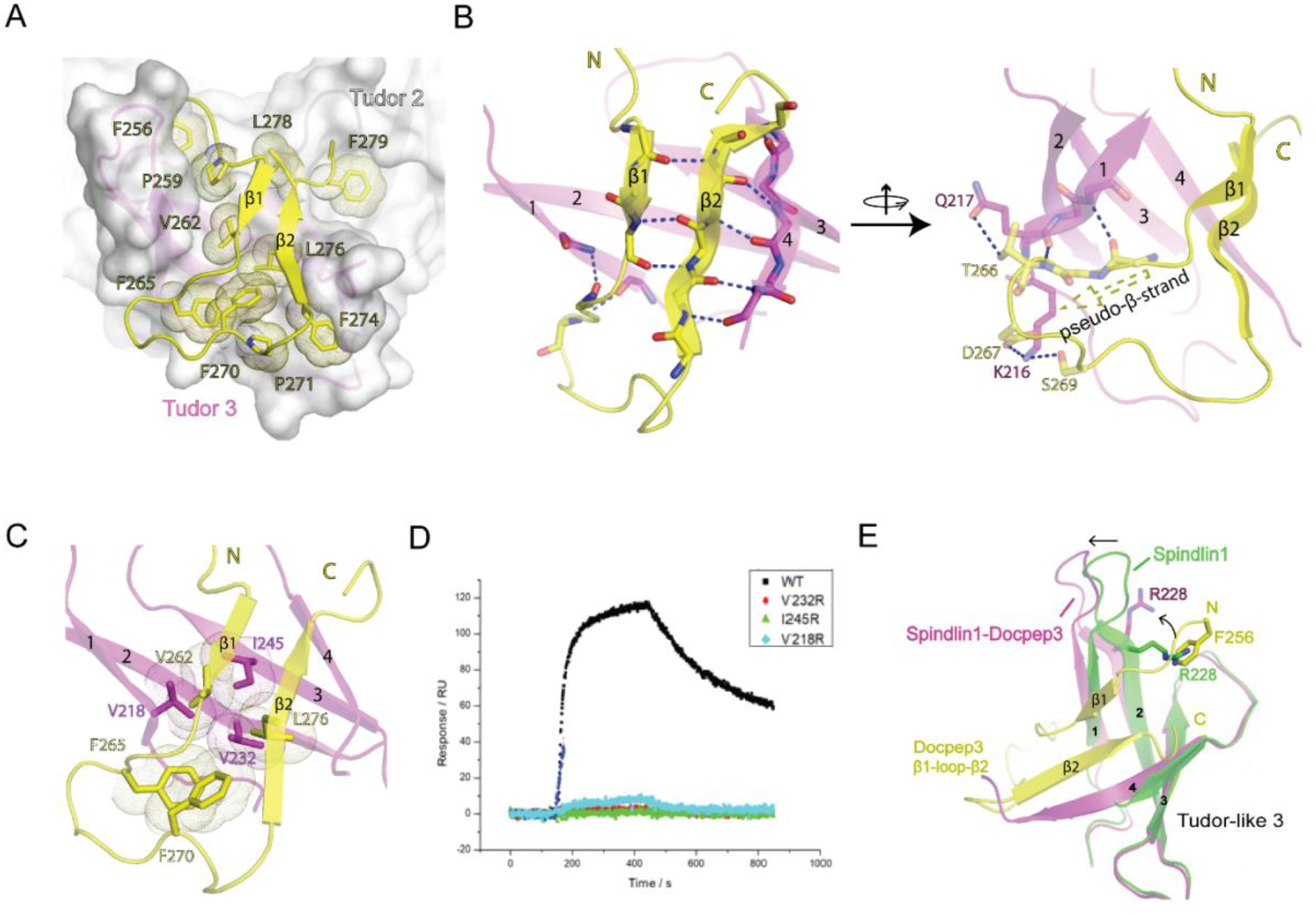
Details of Spindlin1 bound to DOCpep3 and mutagenesis studies. **(A)** Hydrophobic interface between Spindlin1 (surface view in white) and DOCpep3 (ribbon view in yellow) with hydrophobic residues showing dots. **(B)** (left) β-sheet formed by β1, β2 strands (yellow) of DOCpep3 and β4 strand (magenta) of Spindlin1 Tudor 3. (right) A pseudo β-sheet and several hydrogen bonds formed between β1 of Spindlin1 Tudor 3 and DOCpep3. (Blue dashes) direct hydrogen bonds or salt bridges. **(C)** Hydrophobic core formed by Spindlin1 V218, V232, I245 (shown as stick in magenta) and DOCpep3 V262, L276 (shown as stick in yellow). **(D)** SPR binding signals of WT and mutant Spindlin1 flowing immobilized DOCpep3. **(E)** Structural alignment of Spindlin1 Tudor 3 (green) (PDB code: 4MZF) and Spindlin1 Tudor 3 (magenta) bound to DOCpep3 (yellow).

### Two K/R-rich motifs bind to the acidic surface of Spindlin1 Tudor 2

Spindlin1 is characteristic of an acidic surface especially across its Tudor 2, which well meets a need for Tudor 2 in recognizing basic histone tails as well as other K/R-rich peptide such as TCF4 (10,13,15). We then searched for histone like K/R-rich regions of SPIN•DOC and identified two motifs, DOCpep1 (187-195) and DOCpep2 (228-239), which are located N-terminal to DOCpep3 (256-281). To confirm an interaction between each motif and Spindlin1, we synthesized the corresponding peptides and performed isothermal titration calorimetry (ITC) studies. We were able to measure micromolar binding affinities with *K*_d_ values of 78 μM between DOCpep1 and Spindlin1_50-262_ and 31 μM between DOCpep2 and Spindlin1_50-262_ (Figure 3A and B). When the whole K/R-rich region of SPIN•DOC encompassing N184 to L240 was used for titration, an enhanced affinity of 7.6 μM was measured with a calculated binding stoichiometry of 0.5, suggesting two molecules of Spindlin1 bind to one molecule of SPIN•DOC_184-240_ harboring DOCpep1 and DOCpep2 (Figure 3C). The observed 4 to 10-fold binding enhancement suggests that the whole K/R-rich region could associate with Spindlin1 more efficiently likely due to multivalent engagement.

**Figure 3.**
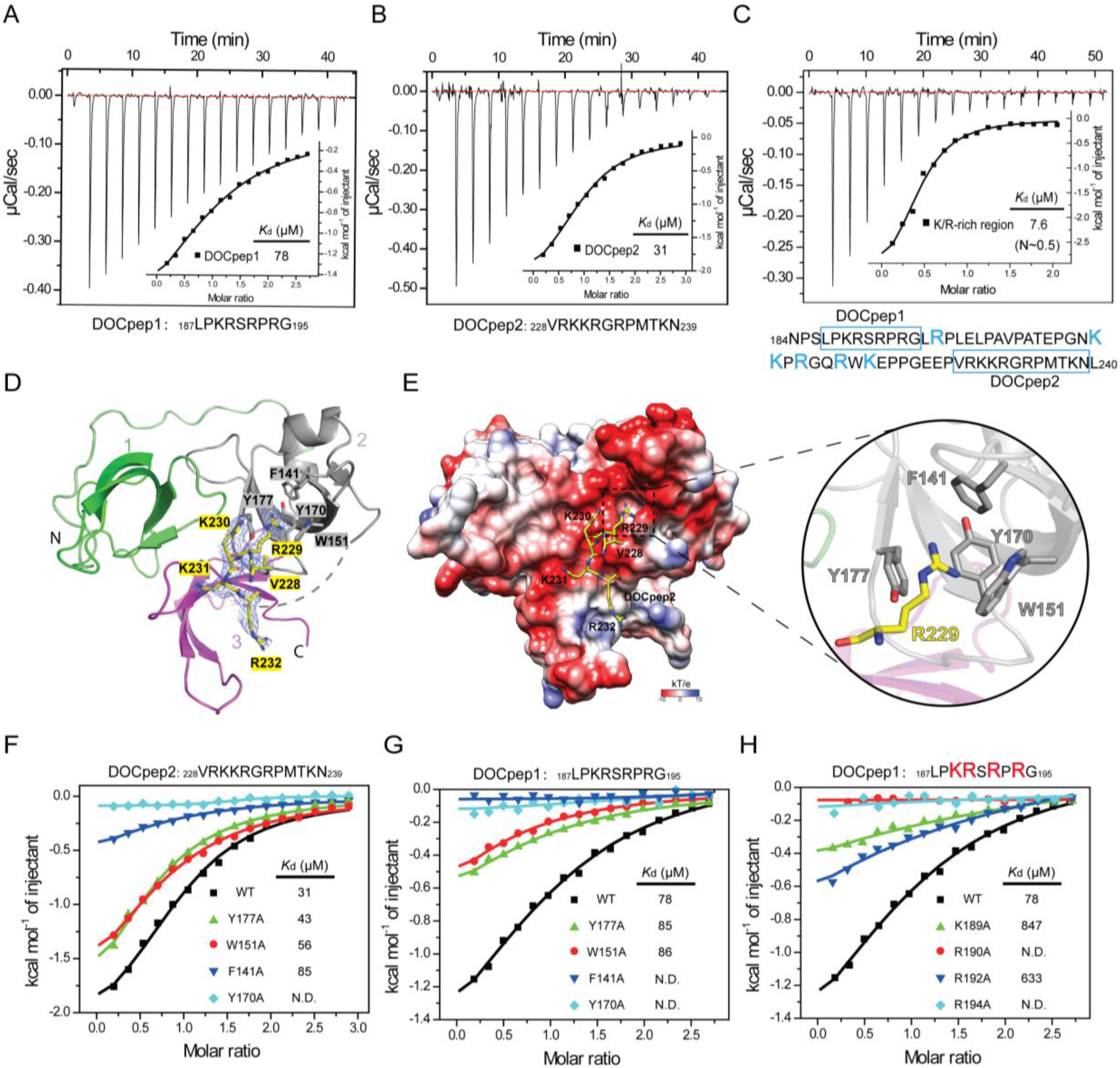
SPIN•DOC K/R-rich region directly binds Spindlin1. **(A, B, C)** ITC titration and fitting curves of Spindlin1 with DOCpep1 (A), DOCpep2 (B) and K/R-rich region (C). **(D)** Overall structure of Spindlin1 bound to DOCpep2 in ribbon view. Based on the electron density, DOCpep2 228-232 is modeled and depicted as yellow sticks. Blue mesh, 2F_o_ - F_c_ omit map around the DOCpep2 contoured at the 0.6σ level. **(E)** (left) Electrostatic potential surface view of Spindlin1-DOCpep2 complex. DOCpep2 is depicted as yellow sticks. Electrostatic potential is expressed as a spectrum ranging from −10 kT/e (red) to +10 kT/e (blue). DOCpep2 is excluded for electrostatic potential calculation. (right) Close-up view of DOCpep2 R229 encapsulated by aromatic cage in Spindlin1’s Tudor 2. **(F, G)** ITC fitting curves of Spindlin1 aromatic pocket mutants in Tudor 2 titrated by DOCpep2 (F) and DOCpep1 (G). **(H)** ITC fitting curves of Spindlin1 titrated by DOCpep1 and its mutants including K189A, R190A, R192A, R194A peptides.

To explore the molecular basis underlying DOCpep1 and DOCpep2 recognition by Spindlin1, we performed crystallization screening and successfully crystallized the DOCpep2-Spindlin1_50-262_ complex and solved its structure at 2.7 Å (Table 1). We were able to model residues V228-R232 of the DOCpep2 peptide according to the electron densities (Figure 3D). In the complex structure, DOCpep2 binds to the acidic surface of Tudor 2 around its histone reader pocket (Figure 3E). Notably, residue R229 is inserted into the aromatic cage formed by residues F141, W151, Y170, and Y177 of Tudor 2, and is stabilized by cation-π interactions (Figure 3E). To validate contributions of these aromatic cage residues in DOCpep2 binding, we generated alanine mutants and performed binding studies. ITC titration revealed that Y177A, W151A, and F141A displayed 1.5-fold to 3-fold binding reduction, while Y170A completely lost binding for DOCpep2, stressing a critical role of the aromatic cage in DOCpep2-Spindlin1 interaction (Figure 3F).

Given the similarity between DOCpep1 and DOCpep2, it is very likely that DOCpep1 binds the same surface of Spindlin1 Tudor 2. In support, the aromatic cage mutants affected DOCpep1-Spindlin1 interaction, with F141A and Y170A totally lost binding (Figure 3G). We also performed alanine mutation of the K/R residues of DOCpep1. Binding studies showed that K189A and R192A led to 9 and 11 fold binding reduction, respectively, while R190A and R194A abolished binding (Figure 3H). Collectively, these data suggest that DOCpep1 and DOCpep2 competitively bind to the same acidic surface around the aromatic cage of Tudor 2. Since the same surface is also involved in histone readout by Spindlin1, our structure and binding studies suggest that SPIN•DOC may inhibit the histone reader activity of Spindlin1 through masking its histone binding surface by its K/R-rich motifs.

### SPIN•DOC attenuates Spindlin1 binding with histone H3 and TCF4 peptides

There is one high affinity hydrophobic motif and two low affinity K/R-rich motifs within SPIN•DOC responsible for Spindlin1 binding. To explore concurrent Spindlin1 engagement by all three motifs, we first performed thermal shift assays comparing Spindlin1_50-262_, Spindlin1_50-262_ bound to DOCpep3, and Spindlin1_50-262_ bound to DOCpep(1-3). As shown in Figure 4A, the melting temperature (Tm) of Spindlin1_50-262_ alone is 49.5 °C, and the Tm value is significantly increased to 59 °C when Spindlin1_50-262_ is bound to DOCpep3. Remarkably, the thermo-stability of Spindlin1_50-262_ is further enhanced by inclusion of additional K/R-rich DOCpep 1 and 2 motifs as reflected by 4.5 °C increase in Tm, stressing a positive role of K/R-rich region in SPIN•DOC-Spindlin1 engagement.

**Figure 4.**
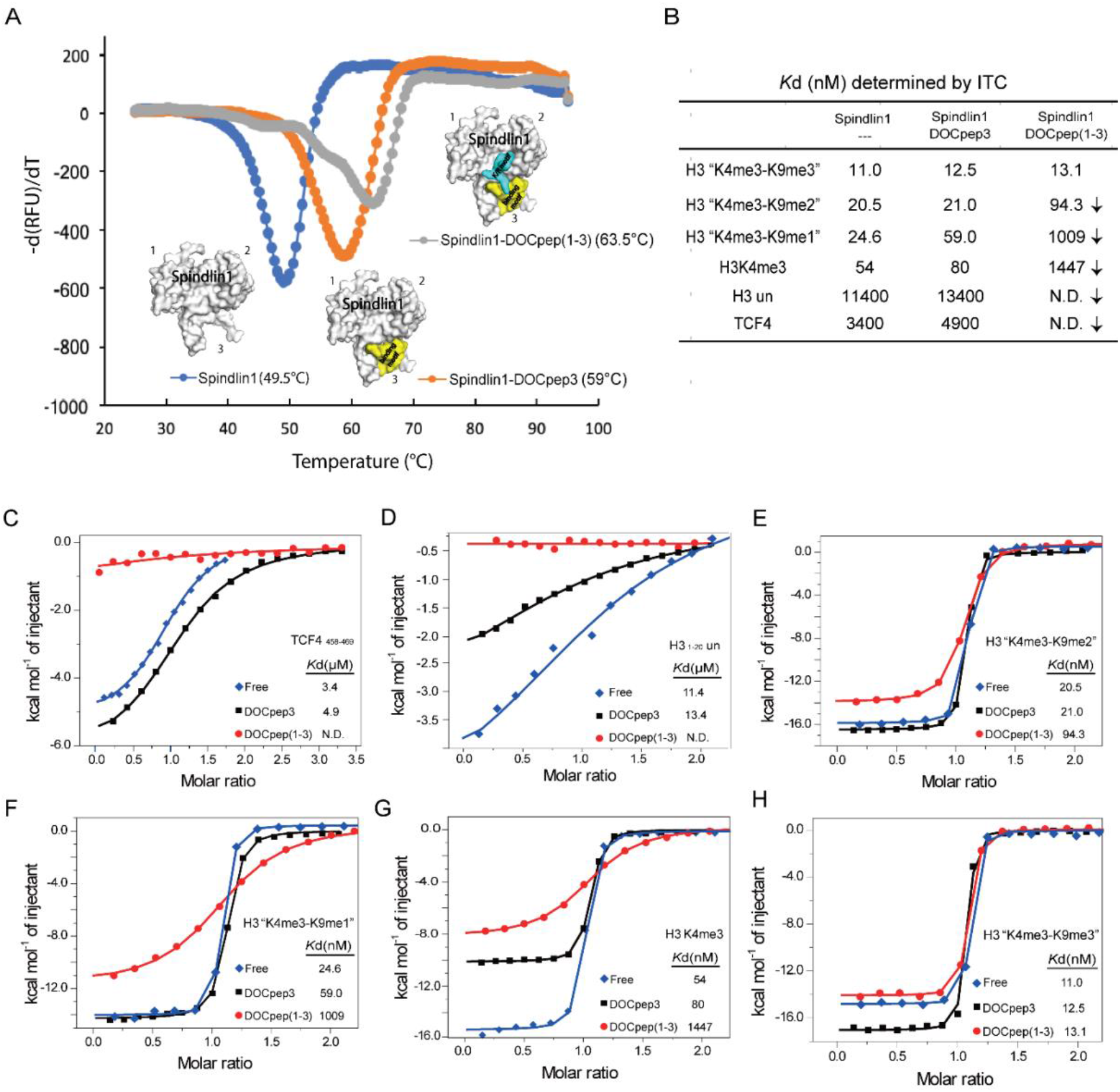
SPIN•DOC enhances Spindlin1’s thermal stability and blocks Spindlin1’s affinity for histone H3 and TCF4 peptides. **(A)** TSA melting curves of Spindlin1_50-262_ alone, Spindlin1_50-262_ bound to SPIN•DOC_256-281_ (DOCpep3), and Spindlin1_50-262_ bound to SPIN•DOC_188-281_ (DOCpep 1-3). **(B)** Summary of *K*_d_ values determined by ITC. The downward arrow indicates a decrease in binding affinity. **(C-G)** ITC fitting curves of free Spindlin1, Spindlin1-DOCpep3 and Spindlin1-DOCpep (1-3) complexes titrated by TCF4458-469 peptide (C), H31-20 un (D), H31-15K4me3K9me2 (E), H31-15K4me3K9me1 (F), H31-20K4me3 (G) and H31-15K4me3K9me3 (H) respectively.

We next evaluated the inhibitory effects of SPIN•DOC on histone H3 and TCF4 peptide binding by Spindlin1. First, the tight engagement with DOCpep3 alone by Spindlin1 is compatible with histone or TCF4 peptide binding, as reflected by similar binding *K*_d_ values to those measured for free Spindlin1 (Figure 4B). This is consisted with the fact that different Tudor domains are involved in DOCpep3 and H3/TCF4 peptides binding. Intriguingly, we observed inhibitory effects to various extent when a longer SPIN•DOC frame encompassing all three DOCpep motifs was used for binding assays. For those low affinity ligands, such as unmodified histone H3 and TCF peptides, their binding with Spindlin1 is fully disrupted by the presence of DOCpep(1-3) (Figure 4C and D). The binding affinities are decreased by ~4.5-fold for H3“K4me3-K9me2” (Figure 4E), ~17-fold for H3“K4me3-K9me1” (Figure 4F), and ~18-fold for H3K4me3 (Figure 4G), respectively. By contrast, Spindlin1’s affinity for H3“K4me3-K9me3” peptide is largely unaffected by the binding of DOCpep(1-3) (Figure 4H). This suggests that an extended SPIN•DOC fragment encompassing DOCpep(1-3) inhibits histone or TCF4 binding by Spindin1 due to introduced competition. Importantly, this inhibitory effect is more pronounced for weaker binding targets but not for strong ones such as H3 “K4me3-K9me3”, suggesting a role of SPIN•DOC in modulating chromatin targeting of Spindlin1 from low affinity to high affinity sites.

### SPIN•DOC down-regulates Spindlin1 target genes via multivalent binding manner

To further validate interaction motifs between Spindlin1 and SPIN•DOC in a cellular context, we performed co-immunoprecipitation (co-IP) using ectopically expressing FLAG-Spindlin1 and GFP-SPIN•DOC with different segments deleted in HEK293T cells. As illustrated in Figure 5A, SPIN•DOC interacts strongly with Spindlin1, while SPIN•DOC with DOCpep3 deleted completely lose the affinity for Spindlin1. Indeed, the fragment DOCpep3 only could interact with Spindlin1 efficiently. The removal of K/R-rich region of SPIN•DOC does not affect its binding to Spindlin1, suggesting DOCpep3 but not K/R-rich region of SPIN•DOC is critical for stable Spindlin1 engagement (Figure 5B).

**Figure 5.**
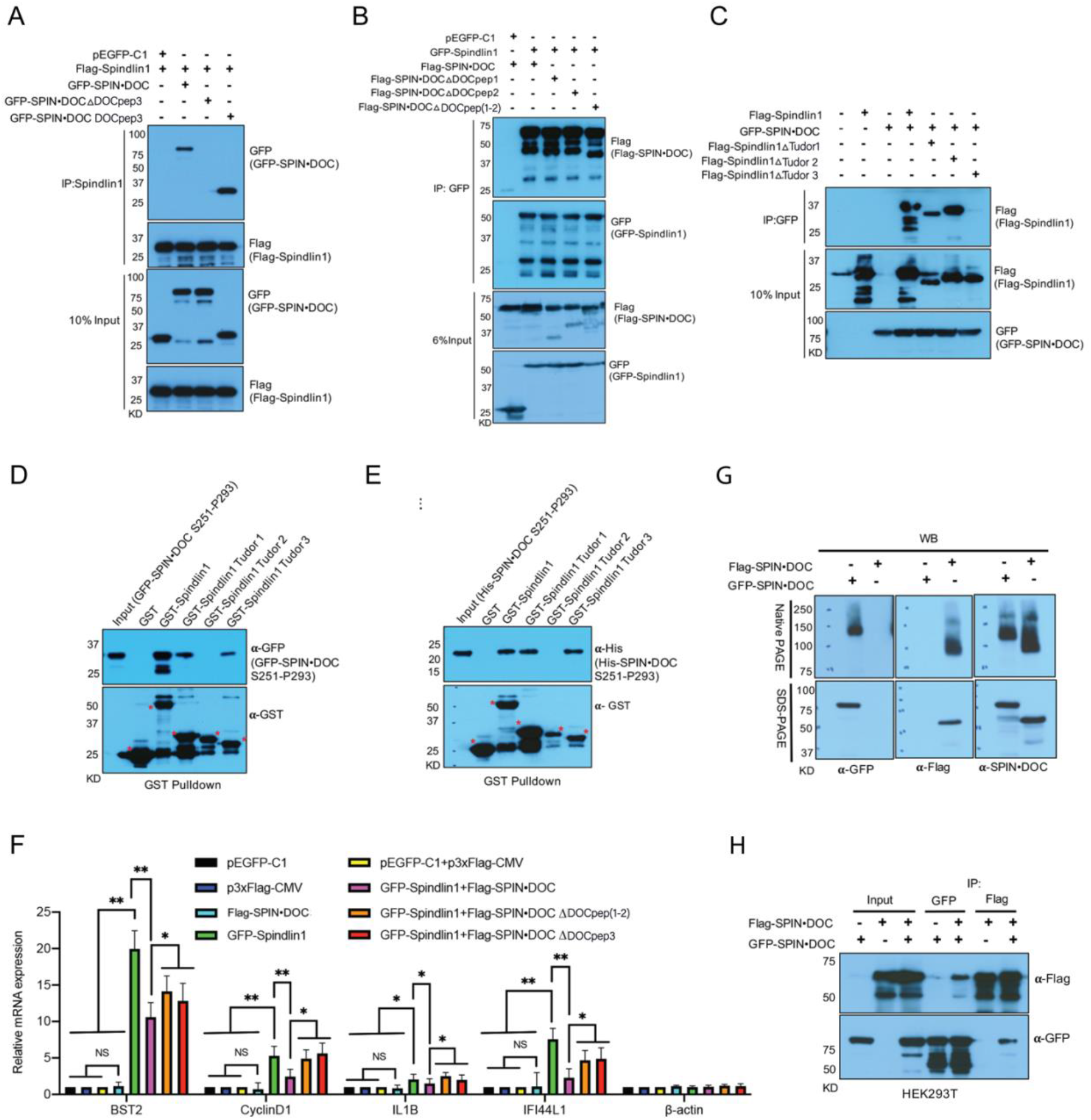
SPIN•DOC attenuates Spindlin1’s transcriptional coactivator activity via a multivalent manner. **(A)** The whole cell lysates from HEK293T cell line co-transfected with Flag-Spindlin1 and GFP-SPIN•DOC wild type and mutants were immunoprecipitated by Spindlin1 antibody and blotted with GFP and Flag antibodies. pEGFP-C1 was an empty vector co-transfected with Flag-Spindlin1 as a negative control. **(B)** The whole cell lysates from HEK293T cell line co-transfected with GFP-Spindlin1 and Flag-SPIN•DOC wild type and mutants were immunoprecipitated by GFP antibody and blotted with GFP and Flag antibodies. pEGFP-C1 was an empty vector co-transfected with Flag-SPIN•DOC as a negative control. **(C)** The whole cell lysates from HEK293T cell line co-transfected with GFP-SPIN•DOC and Flag-Spindlin1 wild type and mutants were immunoprecipitated by GFP antibody and blotted with GFP and Flag antibodies. pEGFP-C1 was an empty vector co-transfected with Flag-Spindlin1 wild type as a control. **(D-E)** GST fusion proteins GST-Spindlin1, GST-Spindlin1 Tudor 1, GST-Spindlin1 Tudor 2 and GST-Spindlin1 Tudor 3 were incubated with cell lysates from HEK293T cell line transfected with GFP-SPIN•DOC S251-P293 (C) and purified fusion protein His-SPIN•DOC S251-P293 (D), then blotted with GFP, His and GST antibodies. GST as a negative control. * indicated corresponding bands. **(F)** Spindlin1 target genes mRNA levels were analyzed by RT-qPCR in T778 cells transfected or co-transfected with vector controls, GFP-Spindlin1, Flag-SPIN•DOC and Flag-SPIN•DOC mutants for 48 hours. Gene expression was normalized to GAPDH levels and β-actin was a gene expression control. Error bars indicate S.D. * *P* < 0.05, ** *P*< 0.01, two-tailed Student’s t test. **(G)** HEK293T cells were transfected with GFP-SPIN•DOC or Flag-SPIN•DOC. Native proteins were extracted for Native PAGE- and SDS-PAGE-Western blot by anti-GFP, anti-Flag and anti-SPIN•DOC. **(H)** HEK293T cells were co-transfected with GFP-SPIN•DOC and Flag-SPIN•DOC, then performed mutual GFP and Flag IPs and analyzed by Western blot.

In the meantime, in order to verify the interaction regions of Spindlin1, GFP-SPIN•DOC and Flag-Spindlin1 with different Tudor domain deleted were expressed in HEK293T cells. The removal of Spindlin1 Tudor 3 caused Spindlin1 to fail to interact with SPIN•DOC. Interestingly, we found that interaction between SPIN•DOC and Tudor1-deleted Spindlin1 also became much weaker compared to the wild type Spindlin1, suggesting Tudor1 may also be involved in SPIN•DOC-Spindlin1 interaction (Figure 5C). To test this possibility, we expressed GFP fused SPIN•DOC_251-293_ in HEK293T cells and then used the lysates for a GST pulldown with the three isolated Tudor domains of Spindlin1 (Figure 5D). Indeed, both Tudor 1 and Tudor 3 are able to pulldown GFP-SPIN•DOC_251-293_. Next, to address whether these interactions are direct, we performed the same pulldown using recombinant His-SPIN•DOC_251-293_. Again, we see an interaction between SPIN•DOC_251-293_ and Spindlin1 Tudor 1 and 3 (Figure 5E). Thus, the SPIN•DOC_251-293_ region can interact with two Tudor domains, which may have interesting regulatory consequences that will be addressed in the discussion section.

Previous studies have shown that SPIN•DOC could inhibit Spindlin1’s transcriptional coactivator activity, significantly reducing the expression level of Spindlin1 target genes (28,29). In order to explore the effects of DOCpep(1-3) on transcriptional coactivator activity of Spindlin1, we performed RT-qPCR analysis to monitor the transcription level of Spindlin1 targeting genes in T778 liposarcoma cells with SPIN•DOC WT or mutants expressed. We found that full-length SPIN•DOC could efficiently inhibit the transcription of Spindlin1 targeting genes, including BST2, CyclinD1, IL1B, and IFI44L1. By contrast, when SPIN•DOC with DOCpep3 or K/R-rich region deleted was expressed, the transcription level of these Spindlin1 targeting genes was compensated to some extent (Figure 5F). This suggests that although DOCpep3 dominants binding of SPIN•DOC to Spindlin1, both DOCpep3 and K/R-rich region (DOCpep1 and 2) are important for SPIN•DOC to attenuate Spindlin1’s transcriptional coactivator activity.

## DISCUSSION

Histone PTMs readout and its precise regulation play important roles in many essential DNA-templated cellular processes, such as transcription, replication, and mitosis. Spindlin1 has been characterized as a multifaceted reader that recognizes histone marks such as H3K4me3, H3K9me3, H4K20me3, H3“K4me3-R8me2a” and H3“K4me3-K9me3/2” depending on the upstream signal and methylation events (7,9,10,13,15). Furthermore, Spindlin1 can also associate with other non-histone partners in different cellular context, such as TCF4 and MAZ transcriptional factors in signaling pathways, the HBx viral protein essential for HBV infection, and SPIN•DOC investigated here (12–14,28,29,32). Overexpression of Spindlin1 leads to upregulated WNT and RET signaling, perturbations of cell cycle and chromosomal instability, as well as tumorigenesis (12, 14, 17, 18, 20, 21, 23–27). So there are growing interests in detecting the regulatory mechanism of Spindlin1, which helps develop target-based inhibitors (33–38). In this study, our biochemical and structural studies established the engagement mode between SPIN•DOC and Spindlin1, which shed mechanistic insights into an inhibitory function of SPIN•DOC against Spindlin1’s transcriptional coactivator activity.

In the SPIN•DOC-Spindlin1 complex, SPIN•DOC is anchored to Spindlin1’s Tudor 3 with a strong binding affinity through a hydrophobic DOCpep3 motif. Besides, a K/R-rich region of SPIN•DOC binds to the acidic surface of Spindlin1 Tudor 2 with weaker affinities. As the K/R-rich region (184-240) and DOCpep3 (256-281) are neighboring motifs, it is conceivable that the high affinity engagement involving DOCpep3 shall bring both DOCpep1 and DOCpep2 motifs spatially proximal to Tudor 2 surface and thus promote their interaction due to increased local concentration, competitively compromising Spindlin1’s association with histone H3K4me3 and TCF4. It is notable that the inhibitory effect of SPIN•DOC against Spindlin1 is more pronounced for weaker binding targets, but not for high affinity ones, such as H3“K4me3-K9me3”. As a result, SPIN•DOC may attenuate Spindlin1’s transcriptional coactivator activity at lower affinity sites and modulate its re-distribution in different chromatin context. In support, our gene expression analysis revealed that SPIN•DOC inhibited expression of Spindlin1 target genes, such as CyclinD1 in WNT signaling pathway, in both DOCpep(1-2) and DOCpep3-dependent manner.

Though lacking structural evidence, our co-IP-based mapping studies suggested a low affinity interaction between Spindlin1 Tudor 1 and SPIN•DOC_251-293_ comprising the DOCpep3 motif. Roles of Tudor 1 as well as Tudor 2 in SPIN•DOC interaction has also been suggested by a serial capture affinity purification-coupled cross-linking mass spectrometry (SCAP-XL) study (39). Of note, an interaction between DOCpep3 and Tudor 3 was not identified by the SCAP-XL strategy likely due to lack of lysine residue in the DOCpep3 motif. Collectively, considering all three Tudors may contribute to SPIN•DOC-Spindlin1 engagement, this raises the possibility that posttranslational modification of either Spindlin1 or SPIN•DOC could control which Tudor mediates this interaction. Alternatively, if SPIN•DOC is able to homodimerize then it may be able to block both Tudor 1 and Tudor 3 functions. Indeed, we investigated the ability of SPIN•DOC to homodimerize, and it does. We demonstrate this homodimerization property by both native gel experiments (Figure 5G), and by coimmunoprecipitation using different tags (Figure 5H). Moreover, dimer formation of SPIN•DOC was also supported by the gel filtration profile of co-expressed full-length SPIN•DOC-Spindlin1 complex (Figure 1A, right).

Based on our study, we proposed a multivalent engagement model for SPIN•DOC-Spindlin1 interaction. First, SPIN•DOC is stably anchored to Spindlin1 Tudor 3 with a super high binding affinity via DOCpep3. Meanwhile, other low affinity engagement regions, especially those K/R-rich motifs, of SPIN•DOC associate with Spindlin1’s acidic surface of Tudor 2 and likely Tudor 1 in a dynamic manner, thus masking the histone binding surface of Spindlin1 to attenuate its transcriptional coactivator activity. In fact, it’s not the first case that an interacting protein may block the histone reader activity of a Tudor domain. 53BP1 is a multi-functional double-strand break (DSB) repair protein and is recruited to DSBs via its tandem Tudor domain recognizing histone H4K20me2. Recently, a previously uncharacterized protein, TIRR (Tudor Interacting Repair Regulator) was demonstrated to directly bind the tandem Tudor domain and mask its H4K20me2 binding motif. Upon DNA damage, ATM phosphorylates 53BP1 and recruits RAP1-ineractiong factor 1 (RIF1) to dissociate the 53BP1-TIRR complex (40). The disassembly of SPIN•DOC-Spindlin1 complex may be similarly regulated in order to recover the coactivator activity of Spindlin1, an intriguing mechanism yet to be established in future studies.

During preparation of this manuscript, a ternary complex structure of Spindlin1 bound to both DOCpep3 and H3“K4me3-K9me3” peptide was published (41). This work reported a transcriptional activating function of the SPIN•DOC-Spindlin1 complex through a proposed HP1-displacement mechanism. In fact, the published structural and binding studies are consistent with our current and previous results, in which SPIN•DOC-Spindlin1 engagement is fully compatible with high affinity readout of the H3 “K4me3-K9me3” mark (7). That said, the chromatin-targeting consequence of SPIN•DOC-Spindlin1 complex is determined by the occurrence of upstream marks and whether their affinity is low or high. Conceivably, the creation of low abundance H3 “K4me3-K9me3/2” bivalent mark and its high affinity readout by SPIN•DOC-Spindlin1 may be critical to trigger derepression of H3K9me3/2-repressed genes such as rDNA repeats. In this study, we revealed additional regulatory mechanism that supports an inhibitory function of SPIN•DOC through affecting chromatin association of Spindlin1 with actively transcribed genes that are co-regulated by weaker yet more prevalent recognition events involving H3K4me3 and TCF4 among other targets (28,29).

## ACCESSION NUMBERS

Atomic coordinates and structure factors for the reported complex structures of Spindlin1-DOCpep3 and Spindlin1-DOCpep2 have been deposited with the Protein Data bank under accession number 7E9M and 7EA1 respectively.

## ACKNOWLEDGEMENT

We thank the staff members at beamline BL17U of the Shanghai Synchrotron Radiation Facility for their assistance in data collection.

## REFERENCE

1. Zhao, S., Zhang, X. and Li, H. (2018) Beyond histone acetylation-writing and erasing histone acylations. Curr Opin Struct Biol, 53, 169–177.

2. Musselman, C.A., Lalonde, M.E., Cote, J. and Kutateladze, T.G. (2012) Perceiving the epigenetic landscape through histone readers. Nat Struct Mol Biol, 19, 1218–1227.

3. Zhang, D., Tang, Z., Huang, H., Zhou, G., Cui, C., Weng, Y., Liu, W., Kim, S., Lee, S., Perez-Neut, M. et al. (2019) Metabolic regulation of gene expression by histone lactylation. Nature, 574, 575–580.

4. Farrelly, L.A., Thompson, R.E., Zhao, S., Lepack, A.E., Lyu, Y., Bhanu, N.V., Zhang, B.C., Loh, Y.H.E., Ramakrishnan, A., Vadodaria, K.C. et al. (2019) Histone serotonylation is a permissive modification that enhances TFIID binding to H3K4me3. Nature, 567, 535–+.

5. Ren, X., Zhou, Y., Xue, Z., Hao, N., Li, Y., Guo, X., Wang, D., Shi, X. and Li, H. (2021) Histone benzoylation serves as an epigenetic mark for DPF and YEATS family proteins. Nucleic Acids Res, 49, 114–126.

6. Greer, E.L. and Shi, Y. (2012) Histone methylation: a dynamic mark in health, disease and inheritance. Nat Rev Genet, 13, 343–357.

7. Zhao, F., Liu, Y., Su, X., Lee, J.E., Song, Y., Wang, D., Ge, K., Gao, J., Zhang, M.Q. and Li, H. (2020) Molecular basis for histone H3 “K4me3-K9me3/2” methylation pattern readout by Spindlin1. J Biol Chem, 295, 16877–16887.

8. Zhao, Q., Qin, L., Jiang, F., Wu, B., Yue, W., Xu, F., Rong, Z., Yuan, H., Xie, X., Gao, Y. et al. (2007) Structure of human spindlin1. Tandem tudor-like domains for cell cycle regulation. J Biol Chem, 282, 647–656.

9. Wang, W., Chen, Z., Mao, Z., Zhang, H., Ding, X., Chen, S., Zhang, X., Xu, R. and Zhu, B. (2011) Nucleolar protein Spindlin1 recognizes H3K4 methylation and stimulates the expression of rRNA genes. EMBO Rep, 12, 1160–1166.

10. Yang, N., Wang, W., Wang, Y., Wang, M., Zhao, Q., Rao, Z., Zhu, B. and Xu, R.M. (2012) Distinct mode of methylated lysine-4 of histone H3 recognition by tandem tudor-like domains of Spindlin1. Proc Natl Acad Sci U S A, 109, 17954–17959.

11. Zhang, X., Zhu, G., Su, X., Li, H. and Wu, W. (2018) Nucleolar localization signal and histone methylation reader function is required for SPIN1 to promote rRNA gene expression. Biochem Biophys Res Commun, 505, 325–332.

12. Franz, H., Greschik, H., Willmann, D., Ozretic, L., Jilg, C.A., Wardelmann, E., Jung, M., Buettner, R. and Schule, R. (2015) The histone code reader SPIN1 controls RET signaling in liposarcoma. Oncotarget, 6, 4773–4789.

13. Su, X., Zhu, G., Ding, X., Lee, S.Y., Dou, Y., Zhu, B., Wu, W. and Li, H. (2014) Molecular basis underlying histone H3 lysine-arginine methylation pattern readout by Spin/Ssty repeats of Spindlin1. Genes Dev, 28, 622–636.

14. Wang, J.X., Zeng, Q., Chen, L., Du, J.C., Yan, X.L., Yuan, H.F., Zhai, C., Zhou, J.N., Jia, Y.L., Yue, W. et al. (2012) SPINDLIN1 Promotes Cancer Cell Proliferation through Activation of WNT/TCF-4 Signaling. Molecular Cancer Research, 10, 326–335.

15. Wang, C., Zhan, L., Wu, M., Ma, R., Yao, J., Xiong, Y., Pan, Y., Guan, S., Zhang, X. and Zang, J. (2018) Spindlin-1 recognizes methylations of K20 and R23 of histone H4 tail. FEBS Lett, 592, 4098–4110.

16. Choi, J.W., Zhao, M.H., Liang, S., Guo, J., Lin, Z.L., Li, Y.H., Jo, Y.J., Kim, N.H. and Cui, X.S. (2017) Spindlin 1 is essential for metaphase II stage maintenance and chromosomal stability in porcine oocytes. Mol Hum Reprod, 23, 166–176.

17. Zhang, P., Cong, B., Yuan, H., Chen, L., Lv, Y., Bai, C., Nan, X., Shi, S., Yue, W. and Pei, X. (2008) Overexpression of spindlin1 induces metaphase arrest and chromosomal instability. J Cell Physiol, 217, 400–408.

18. Choi, J.W., Zhou, W., Nie, Z.W., Niu, Y.J., Shin, K.T. and Cui, X.S. (2019) Spindlin1 alters the metaphase to anaphase transition in meiosis I through regulation of BUB3 expression in porcine oocytes. J Cell Physiol, 234, 8963–8974.

19. Oh, B., Hwang, S.Y., Solter, D. and Knowles, B.B. (1997) Spindlin, a major maternal transcript expressed in the mouse during the transition from oocyte to embryo. Development, 124, 493–503.

20. Sun, M., Li, Z. and Gui, J.F. (2010) Dynamic distribution of spindlin in nucleoli, nucleoplasm and spindle from primary oocytes to mature eggs and its critical function for oocyte-to-embryo transition in gibel carp. J Exp Zool A Ecol Genet Physiol, 313, 461–473.

21. Yuan, H., Zhang, P., Qin, L., Chen, L., Shi, S., Lu, Y., Yan, F., Bai, C., Nan, X., Liu, D. et al. (2008) Overexpression of SPINDLIN1 induces cellular senescence, multinucleation and apoptosis. Gene, 410, 67–74.

22. Wang, J.X., Zeng, Q., Chen, L., Du, J.C., Yan, X.L., Yuan, H.F., Zhai, C., Zhou, J.N., Jia, Y.L., Yue, W. et al. (2012) SPINDLIN1 promotes cancer cell proliferation through activation of WNT/TCF-4 signaling. Mol Cancer Res, 10, 326–335.

23. Fang, Z., Cao, B., Liao, J.M., Deng, J., Plummer, K.D., Liao, P., Liu, T., Zhang, W., Zhang, K., Li, L. et al. (2018) SPIN1 promotes tumorigenesis by blocking the uL18 (universal large ribosomal subunit protein 18)-MDM2-p53 pathway in human cancer. Elife, 7.

24. Gao, Y., Yue, W., Zhang, P., Li, L., Xie, X., Yuan, H., Chen, L., Liu, D., Yan, F. and Pei, X. (2005) Spindlin1, a novel nuclear protein with a role in the transformation of NIH3T3 cells. Biochem Biophys Res Commun, 335, 343–350.

25. Janecki, D.M., Sajek, M., Smialek, M.J., Kotecki, M., Ginter-Matuszewska, B., Kuczynska, B., Spik, A., Kolanowski, T., Kitazawa, R., Kurpisz, M. et al. (2018) SPIN1 is a proto-oncogene and SPIN3 is a tumor suppressor in human seminoma. Oncotarget, 9, 32466–32477.

26. Lv, B.B., Ma, R.R., Chen, X., Zhang, G.H., Song, L., Wang, S.X., Wang, Y.W., Liu, H.T. and Gao, P. (2020) E2F1-activated SPIN1 promotes tumor growth via a MDM2-p21-E2F1 feedback loop in gastric cancer. Mol Oncol.

27. Zhao, M., Bu, Y., Feng, J., Zhang, H., Chen, Y., Yang, G., Liu, Z., Yuan, H., Yuan, Y., Liu, L. et al. (2020) SPIN1 triggers abnormal lipid metabolism and enhances tumor growth in liver cancer. Cancer Lett, 470, 54–63.

28. Bae, N., Gao, M., Li, X., Premkumar, T., Sbardella, G., Chen, J. and Bedford, M.T. (2017) A transcriptional coregulator, SPIN.DOC, attenuates the coactivator activity of Spindlin1. J Biol Chem, 292, 20808–20817.

29. Devi, M.S., Meiguilungpou, R., Sharma, A.L., Anjali, C., Devi, K.M., Singh, L.S. and Singh, T.R. (2019) Spindlin docking protein (SPIN.DOC) interaction with SPIN1 (a histone code reader) regulates Wnt signaling. Biochem Biophys Res Commun, 511, 498–503.

30. Emsley, P. and Cowtan, K. (2004) Coot: model-building tools for molecular graphics. Acta Crystallogr D Biol Crystallogr, 60, 2126–2132.

31. Adams, P.D., Afonine, P.V., Bunkoczi, G., Chen, V.B., Davis, I.W., Echols, N., Headd, J.J., Hung, L.W., Kapral, G.J., Grosse-Kunstleve, R.W. et al. (2010) PHENIX: a comprehensive Python-based system for macromolecular structure solution. Acta Crystallogr D Biol Crystallogr, 66, 213–221.

32. Ducroux, A., Benhenda, S., Riviere, L., Semmes, O.J., Benkirane, M. and Neuveut, C. (2014) The Tudor domain protein Spindlin1 is involved in intrinsic antiviral defense against incoming hepatitis B Virus and herpes simplex virus type 1. PLoS Pathog, 10, e1004343.

33. Bae, N., Viviano, M., Su, X.N., Lv, J., Cheng, D.H., Sagum, C., Castellano, S., Bai, X., Johnson, C., Khalil, M.I. et al. (2017) Developing Spindlin1 small-molecule inhibitors by using protein microarrays. Nat Chem Biol, 13, 750–+.

34. Fagan, V., Johansson, C., Gileadi, C., Monteiro, O., Dunford, J.E., Nibhani, R., Philpott, M., Malzahn, J., Wells, G., Faram, R. et al. (2019) A Chemical Probe for Tudor Domain Protein Spindlin1 to Investigate Chromatin Function. J Med Chem, 62, 9008–9025.

35. Luise, C. and Robaa, D. (2018) Application of Virtual Screening Approaches for the Identification of Small Molecule Inhibitors of the Methyllysine Reader Protein Spindlin1. Methods Mol Biol, 1824, 347–370.

36. Papagiannopoulos, C.I., Theodoroula, N.F., Kyritsis, K.A., Akrivou, M.G., Kosmidou, M., Tsouderou, K., Grigoriadis, N. and Vizirianakis, I.S. (2021) The histone methyltransferase inhibitor A-366 enhances hemoglobin expression in erythroleukemia cells upon co-exposure with chemical inducers in culture. J Biol Res-Thessalon, 28.

37. Robaa, D., Wagner, T., Luise, C., Carlino, L., McMillan, J., Flaig, R., Schule, R., Jung, M. and Sippl, W. (2016) Identification and Structure-Activity Relationship Studies of Small-Molecule Inhibitors of the Methyllysine Reader Protein Spindlin1. Chemmedchem, 11, 2327–2338.

38. Wagner, T., Greschik, H., Burgahn, T., Schmidtkunz, K., Schott, A.K., McMillan, J., Baranauskiene, L., Xiong, Y., Fedorov, O., Jin, J. et al. (2016) Identification of a small-molecule ligand of the epigenetic reader protein Spindlin1 via a versatile screening platform. Nucleic Acids Research, 44.

39. Liu, X., Zhang, Y., Wen, Z., Hao, Y., Banks, C.A.S., Lange, J.J., Slaughter, B.D., Unruh, J.R., Florens, L., Abmayr, S.M. et al. (2020) Driving integrative structural modeling with serial capture affinity purification. Proc Natl Acad Sci U S A, 117, 31861–31870.

40. Drane, P., Brault, M.E., Cui, G., Meghani, K., Chaubey, S., Detappe, A., Parnandi, N., He, Y., Zheng, X.F., Botuyan, M.V. et al. (2017) TIRR regulates 53BP1 by masking its histone methyl-lysine binding function. Nature, 543, 211–216.

41. Du, Y., Yan, Y., Xie, S., Huang, H., Wang, X., Ng, R.K., Zhou, M.M. and Qian, C. (2021) Structural mechanism of bivalent histone H3K4me3K9me3 recognition by the Spindlin1/C11orf84 complex in rRNA transcription activation. Nat Commun, 12, 949.

